# Deciphering the circular RNAs landscape in amyotrophic lateral sclerosis

**DOI:** 10.1101/2025.09.25.677871

**Authors:** Reza Ataei, Javad Amini, Nima Sanadgol, Anne F Simon, Martin L Duennwald

**Affiliations:** Department of Biology, University of Western Ontario, Canada; Products and Medicinal Plants Research Center, North Khorasan University of Medical Sciences, Bojnurd, Iran; Institute of Neuroanatomy, RWTH University Hospital Aachen, Aachen, Germany; Department of Anatomy and Cell Biology, Schulich School of Medicine and Dentistry, University of Western Ontario, Canada

**Keywords:** Amyotrophic lateral sclerosis (ALS), circular RNAs (circRNAs), miRNA sponging, FUS

## Abstract

Amyotrophic lateral sclerosis (ALS) is a fatal neurodegenerative disorder characterized by the progressive loss of motor neurons, with most cases lacking a clear genetic basis. Emerging evidence highlights the involvement of non-coding RNAs, particularly circular RNAs (circRNAs), in disease onset and progression. In this study, we investigated circRNAs implicated in ALS and related motor neuron diseases (MNDs). First, we conducted a systematic review to identify ALS-associated circRNAs, followed by in silico analyses of 15 selected candidates. Our results revealed that hsa_circ_0000099, hsa_circ_0001017, hsa_circ_004846, and hsa_circ_0034880 regulate a high number of ALS-related genes through miRNA sponging. Pathway enrichment analysis indicated that hsa_circ_0000099 is particularly involved in unfolded protein response, oxidative stress, cell cycle regulation, and apoptosis. Protein–RNA interaction analysis further showed that ALS-related circRNAs can sponge 20 RNA-binding proteins, with FMR1, ELAVL1, EIF4A3, and SRSF1 exhibiting the highest number of interactions. Additionally, molecular docking analysis demonstrated that FUS mutations significantly alter its binding affinity to hsa_circ_0000567 and hsa_circ_0060762. RNA-seq data from ALS patients confirmed significant alterations in the expression of host genes of ALS-related circRNAs and hub proteins across affected tissues, including the spinal cord and multiple brain regions. Collectively, these findings support circRNAs as active contributors to ALS pathogenesis. In particular, the host gene of hsa_circ_0000099, which is expressed in the spinal cord and hippocampus, emerges as a potential biomarker for tissue-specific impairment in ALS. Moreover, its regulatory role in key ALS-related cellular pathways underscores its promise as a candidate for biomarker development and therapeutic targeting. Further experimental validation is warranted to confirm its role in ALS pathology.

**Graphical abstract:** 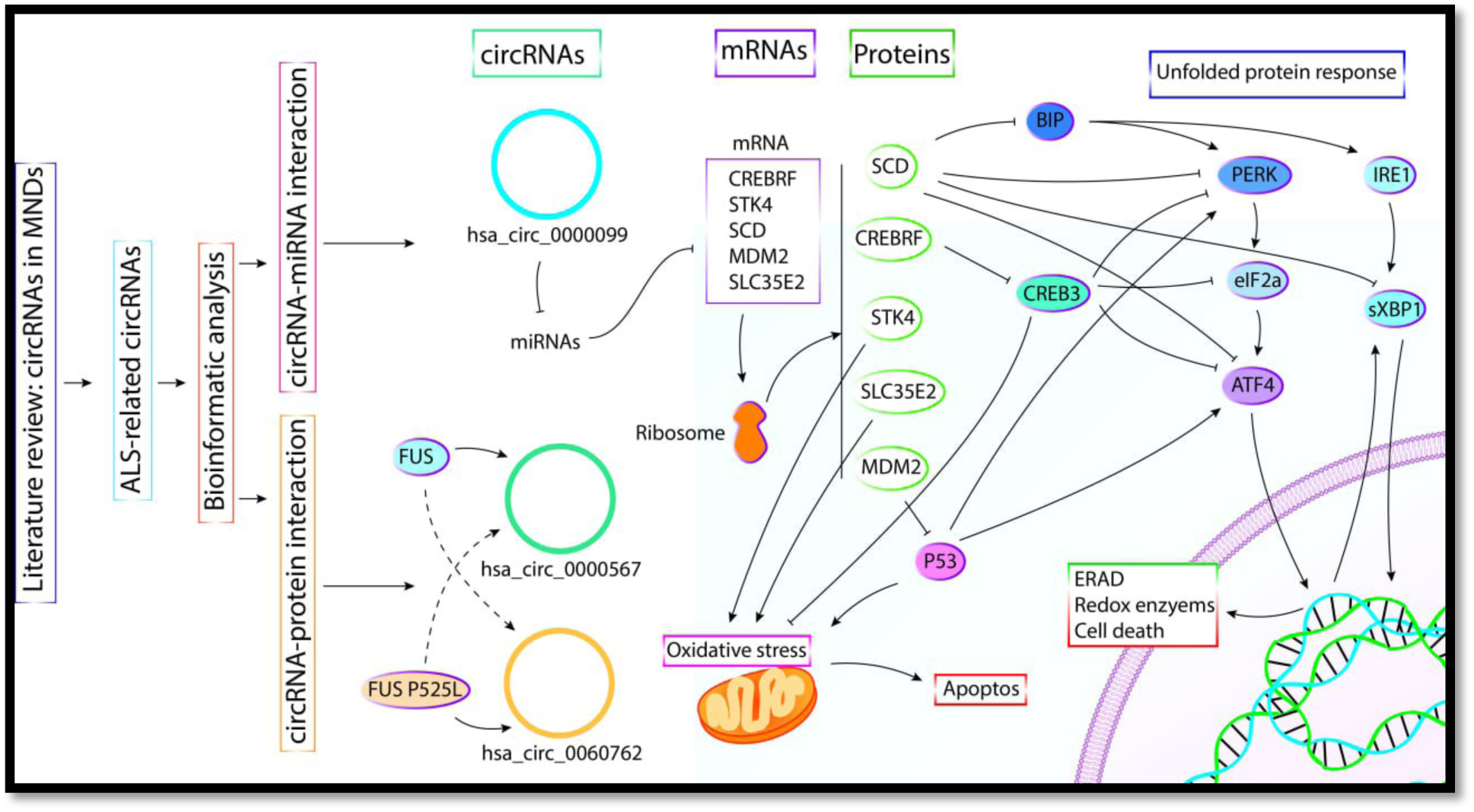

## 1. Introduction

Amyotrophic lateral sclerosis (ALS) results from the dysfunction of upper motor neurons in the motor cortex and lower motor neurons in the ventral horn of the spinal cord, causing muscle weakness without causing significant sensory symptoms or pain (1). The prevalence of ALS, also known as Lou Gehring’s disease, ranges between 4 and 8 cases per 100,000 people, varying across different populations (2). ALS targets motor neurons, leading to gradual loss of these neurons in the brain and spinal cord. The condition typically begins with a localized onset, but as it advances, it extends to other regions of the body, with a survival range of 2 to 5 years following the onset of the disease (3). Sporadic ALS (sALS) represents 90% of all ALS cases, whereas familial ALS (fALS) comprises approximately 10% (4).

A consistent hallmark of ALS pathology is the abnormal aggregation of disease-related proteins. Cytoplasmic inclusions of TAR DNA binding protein (TDP-43/TARDBP) are observed in more than 95% of ALS cases, while other proteins such as superoxide dismutase 1 (SOD1), FUS RNA binding protein (FUS), and dipeptide repeat proteins in C9orf72-SMCR8 complex subunit (C9orf72) carriers also form toxic aggregates, reflecting a failure of proteostasis (5). Mutations in more than 30 distinct genes have been identified as causes of fALS, with SOD1, TARDBP, FUS, and C9orf72 alone accounting for nearly 60% of cases; despite this genetic heterogeneity, these mutations often converge on similar metabolic dysfunctions (4), (6). Disrupted RNA metabolism, excessive accumulation of protein–RNA assemblies, abnormal generation of ribonucleoprotein granules, and defective autophagy-mediated protein clearance are increasingly recognized as key contributors to ALS/FTD development (7). A high number of ALS-associated genes encode RNA-binding proteins that regulate various aspects of RNA metabolism, including mRNA transcription, transport, stabilization, and miRNA production. Emerging evidence indicates that disruption of RNA homeostasis represents a core pathogenic mechanism in ALS (8). For instance, cross-linking immunoprecipitation (CLIP)-Seq analysis identified over 39,000 binding sites of TDP-43 across the mouse transcriptome. In addition, depletion of TDP-43 in the adult mouse brain led to altered splicing of 965 mRNAs, most of which encode proteins involved in synaptic function, highlighting the protein’s essential role in maintaining normal splicing of brain-enriched transcripts. Likewise, in FUS-linked ALS, widespread changes in alternative splicing were observed, resulting in disrupted neuronal gene expression and the generation of thousands of abnormally processed mRNAs (9).

Despite advances in understanding ALS genetics and pathology, clinical research has been unsuccessful in devising effective methods to restore lost motor neurons due to the challenges associated with neuronal regeneration (10). Riluzole, the sole drug authorized for use in Australia, extends median life expectancy by only 2 to 3 months (11). Nevertheless, genetic insights are beginning to inform therapeutic innovation. More recently, genetic insights have paved the way for targeted therapies such as antisense oligonucleotides designed to silence mutant SOD1 (12). (13).

Alongside these protein-coding gene targets, RNA metabolism has emerged as a crucial factor in ALS pathogenesis, drawing attention to the role of non-coding RNAs. Non-coding RNAs play a dynamic role in motor neuron diseases (MNDs). They are highly expressed in the nervous system and contribute significantly to neural development, function, and pathology (14). Non-coding RNAs are categorized based on their length, including microRNAs (miRNAs) and long non-coding RNAs (lncRNAs). LncRNAs can be found inside cells or in the extracellular environment, appearing in either a linear form or as circular RNAs (circRNAs) (15). Unlike their linear counterparts, circRNAs represent a category of either naturally occurring (biogenic) or artificially synthesized closed RNA molecules that lack 5′ and 3′ ends; their covalently closed-loop structure renders them resistant to exonuclease-mediated degradation, granting them greater stability than both lncRNAs and linear mRNAs (16), (17). More than 10% of expressed genes are capable of generating circRNAs. In humans, 5.8–23% of actively transcribed genes contribute to circRNA production. Their expression follows a complex pattern that is tissue-specific, cell-type-specific, or dependent on developmental stages. CircRNAs are particularly prevalent in the brain and during fetal development, with nearly 20% of protein-coding genes producing circRNAs in mammalian brains (17). CircRNAs are highly abundant in neuronal synapses and play a crucial role in regulating the pathological processes associated with neurodegenerative diseases, and they exert their effects through multiple signaling pathways (18). Due to their unique characteristics and tissue-specific expression, circRNAs hold significant potential as biomarkers and vaccine candidates (19).

In this study, we set out to review and analyze the role of circRNAs across the spectrum of MNDs using bioinformatics tools. However, because the majority of available circRNA data were concentrated in ALS, our analysis predominantly emphasizes ALS-related circRNAs while maintaining the original broader scope. We highlight their potential functions and aim to propose key circRNAs as potential candidates for further functional investigation.

## 2. Metabolism of circRNAs

### 2.1. Biogenesis of circRNAs

CircRNA biogenesis depends on the canonical spliceosomal machinery and is specific to certain cell types. In linear splicing, an upstream 5′ splicing site is joined to a downstream 3′ splicing site (Figure 1). Conversely, backsplicing involves looping intron sequences by linking the downstream 5′ splicing site to the upstream 3′ splicing site. The process of backsplicing is influenced by cis-acting elements (like Alu elements), trans-acting splicing factors, and RNA-binding proteins (20). These RNA transcripts exhibited downstream exons joined to upstream exons at unique “back-splice” sites, a phenomenon not accounted for by traditional linear splicing of primary transcripts. These back-splice sites were identified as the circular junctions characteristic of circular RNAs. The majority of circRNAs are exonic, derived from mRNA host genes and utilizing the same exon boundaries as their linear mRNA counterparts (21). Circular RNAs are classified into three types: circular intronic RNAs (ciRNAs), exon-intron circRNAs (EIciRNAs), and exonic circRNAs (ecircRNAs). This categorization is based on their splicing mechanisms and the inclusion of exons and/or introns in their structure (22).

**Figure 1.**
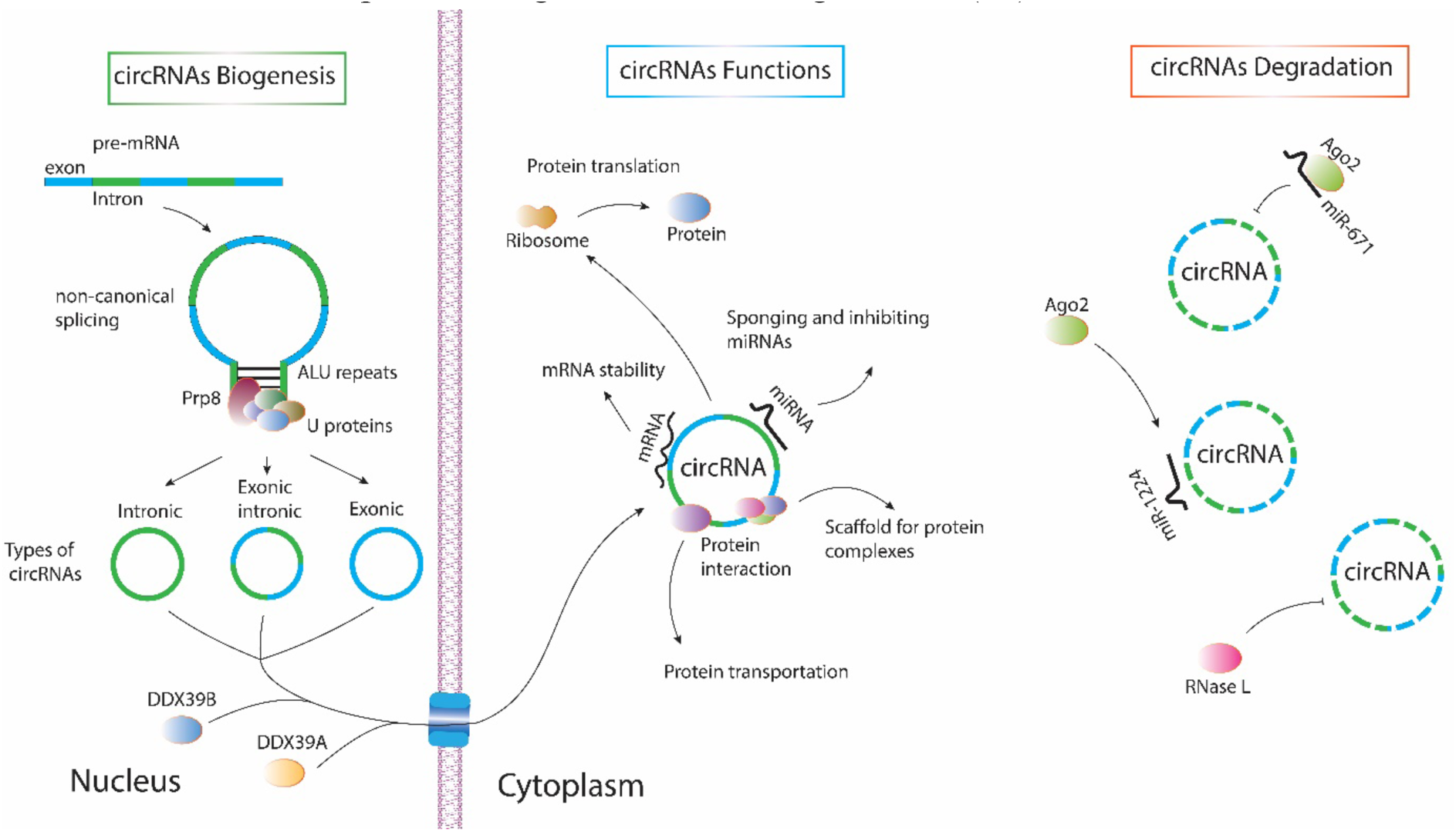
Biogenesis of circRNAs. After transcription, mRNAs are processed by cellular proteins through splicing. Typically, this results in mature mRNAs via canonical splicing. However, when the ends of an mRNA connect, it undergoes non-canonical splicing, producing circRNAs. These circRNAs play multiple roles in the cell, such as miRNA sponging, involvement in translation, and interaction with various proteins.

### 3.2. Regulation of circRNAs formation

Some sequences on mRNAs have an influence in the formation of circRNAs. RNA duplexes frequently originate from repetitive elements, such as primate Alus, which can form inverted repeat Alus (IRAlus). However, some RNA pairs arise from non-repetitive complementary sequences. Additionally, short repeats (∼30 to 40 nucleotides) are adequate to promote circRNA formation in expression vectors (23). The spliceosome is a sizable molecular complex consisting of five small nuclear RNAs (snRNAs) that associate with proteins to form small nuclear ribonucleoproteins (snRNPs). Research indicates that fluctuations in the expression of key spliceosome components significantly influence circRNA production levels. For instance, critical elements such as U2 snRNP (SF3B1 or SF3A1), snRNP-U1-70K, and snRNP-U1-C contribute to 5′ splice site recognition, while the spliceosome’s largest protein component (Prp8) interacts with U5 and U6 snRNAs. When these components become depleted, pre-mRNA splicing is redirected toward circRNA formation, leading to a reduction in linear RNA production (24). RNA-binding proteins (RBPs) play a crucial role in facilitating tissue-specific circRNA formation by interacting with specific motifs within flanking intron sequences. For instance, ALU repeats serve as primary binding sites for adenosine deaminase 1 acting on RNA (ADAR1). ADAR1 suppresses circRNA formation by destabilizing paired elements, such as ALU repeats, through A→I RNA editing in double-stranded RNA (dsRNA) substrates, where adenosines are converted to inosines (25). Transportation of circRNAs from nucleus to cytoplasm mostly facilitate by DExD-Box Helicase 39B (DDX39B) or DExD-Box Helicase 39A (DDX39A) proteins. Despite stability of circRNAs, can be globally degraded by endonuclease, RNase L. Moreover, there are some evidences for role of UPF1 RNA helicase and ATPase (UPF1) and G3BP stress granule assembly factor 1 (G3BP1) in decay of circRNAs (26).

### 2.3. Functions of circRNAs

Several functions have been introduced for circRNAs including 1) sponging miRNAs, 2) protein transportation, 3) transcription regulation, 4) translating to proteins, and 5) stability of mRNAs (27). Circular RNAs possess common binding sites for miRNAs, allowing them to function as miRNA sponges. By capturing and inactivating miRNAs, they regulate the expression of mRNAs targeted by these miRNAs (28). circRNAs exhibit sponging effects on RNA-binding proteins (RBPs), allowing them to bind, isolate, or transport RBPs to specific subcellular locations. Moreover, circRNAs can act as protein scaffolds, facilitating the assembly of various molecules and protein complexes (29). Certain ecircRNAs, like circZNF827 and circACTN4, residing in the nucleus, can stimulate gene transcription by engaging with promoters and attracting transcription regulator proteins to the promoter region. Additionally, ciRNAs, such as ci-ankrd52, and EIciRNAs play a role in transcription regulation by associating with RNA polymerase II within the nucleus of human cells (30). CircRNAs were initially regarded as noncoding RNAs due to the absence of 5′ caps and poly(A) tails, which are crucial for efficient translation. However, numerous studies have revealed that circRNAs possess coding potential. Recent research indicates that the translation of circRNAs relies on internal ribosome entry sites (IRESs). Notably, engineered circRNA expression vectors containing IRESs have been shown to produce functional proteins (31). CircRNAs also serve additional roles, including stabilizing mRNAs and facilitating their transport from the cell body to the extracellular fluid via exosomes (27).

### 2.4. circRNAs degradation

Ago2, a widely expressed Argonaute protein, plays a role in circRNA degradation. miR-671 clears circRNA-CDR1as in an Ago2-dependent manner, while miRNA-1224 regulates circRNA-Filip1l by forming a complex with its precursor, which Ago2 then cleaves, reducing mature circRNA-Filip1l in the spinal nucleus. Unlike miRNA-1224, miR-671 binds almost entirely to circRNA-CDR1as in the nucleus, allowing Ago2 to recognize and degrade it. Moreover, Endonucleases, like RNase H1, contribute to circRNA degradation. Certain circRNAs, such as ci-ankrd52, form stable R-loops with template DNA, but these structures are susceptible to RNase H1 cleavage, leading to their degradation (20). A recent study by Park and colleagues revealed that the m6A reader protein YTHDF2 identifies m6A modifications on circular RNAs and interacts with the adaptor protein HRSP12, which subsequently recruits the RNase P/MRP complex for targeted circRNA degradation (32).

## 3. Material and methods

### 3.1. Searching and scanning the eligible articles

This review was carried out in adherence to the guidelines outlined in the Preferred Reporting Items for Systematic Reviews and Meta-Analyses (PRISMA) (33). An extensive literature search was conducted using three prominent scientific databases—PubMed, Web of Science, and Scopus—to include publications spanning the period from January 2010 to April 2025. Notably, a study focusing on non-coding RNAs, including circRNAs, in ALS was published in December 2021. This emphasizes the innovative nature of these ncRNAs and reflects the increasing scientific attention on their roles. In this study, we used specific MeSH terms, including ALS circRNA, Primary Lateral Sclerosis circRNA, Progressive Muscular Atrophy circRNA, and Spinal Muscular Atrophy circRNA. To minimize bias, the literature search was carried out by three independent authors, and subsequently verified by two additional independent reviewers.

#### 3.1.2. Selection criteria

Given the limited number of studies focusing on circRNAs in MNDs, all relevant investigations addressing circRNAs in this context were included, such as those exploring biomarkers and experimentally validated functional circRNAs. Both in vitro and in vivo studies involving human and animal sources were considered. However, preprint articles and non-English publications were excluded. The literature search yielded a collection of publications that underwent screening by three authors (J.A., R.A.), with each author assigned to one database. The screening process was conducted in two steps. In the first step, titles, abstracts, and keywords were examined to eliminate studies that did not meet the eligibility criteria. At this stage, review articles, conference abstracts, preprints, non-English publications, and unpublished manuscripts were excluded. Following this, duplicate articles were discarded. Subsequently, studies with unrelated titles, such as those focusing on other diseases or conditions, were removed. Lastly, the abstracts of the selected articles were reviewed to verify their alignment with our criteria. In the second step, the full-text articles were evaluated against the inclusion criteria, and the pertinent data were extracted. The selected publications were meticulously evaluated by two authors (J.A. and N.S) according to the established inclusion and exclusion criteria. Adhering to PRISMA guidelines, as depicted in the PRISMA flow diagram (Fig. 2), the search process was conducted transparently and reproducibly.

**Figure 2.**
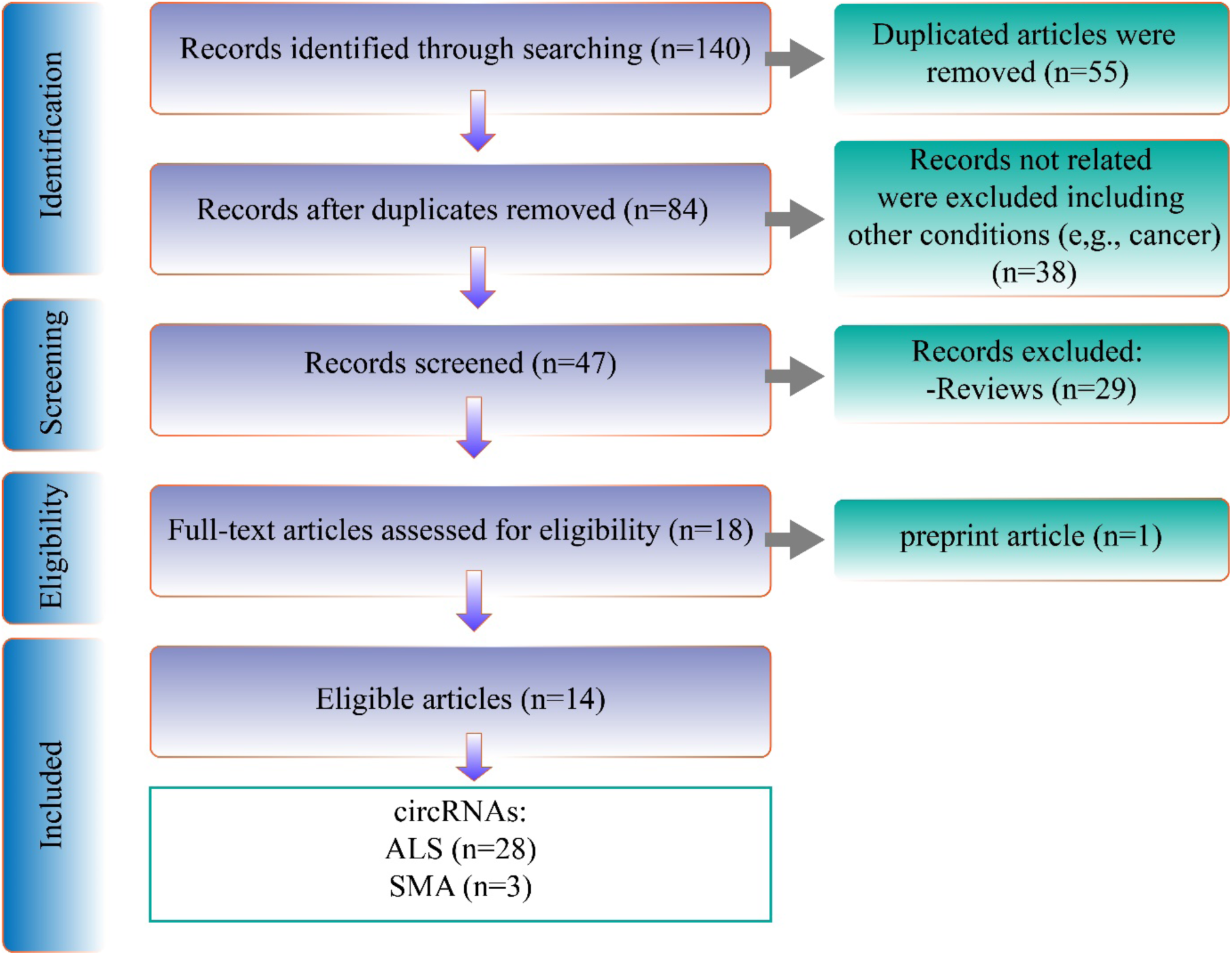
The steps of article selection. After a significant verify 14 articles were selected out of 140 articles. Total 28 circRNAs were found for ALS and 3 circRNAs for SMA.

#### 3.1.3. Data extraction

The data extracted from each selected study included the following components: the author, year of publication, the pathophysiological role of circRNAs (whether pathogenic or protective), the protein or miRNA targets of circRNAs, their downstream effects in MNDs, the study type (human or animal), the investigated tissue or cell type, the circBase code of the circRNAs, and their sequences.

### 3.2. In silico analysis

#### 3.2.1. Extracting basic data of circRNA

We conducted a bioinformatic analysis of circRNAs that have a circBase ID, as the database provides the complete sequence—essential for bioinformatic analysis—along with host genes and other valuable information about circRNAs. Therefore, we proceeded bioinformatic analysis by hsa_circ_0063411, hsa_circ_0088036, hsa_circ_0023919, hsa_circ_0060762, hsa_circ_0000119, hsa_circ_0000567, hsa_circ_0007778, hsa_circ_0000099, hsa_circ_0005171, hsa_circ_0125620, hsa_circ_0001017, hsa_circ_0001439, hsa_circ_0008870, hsa_circ_0034880, and hsa_circ_0004846 circRNAs. Information on circRNAs, including their chromosomal location, associated host gene, and nucleotide sequence, was obtained from the circBase database. CircBase serves as an open-access online platform that curates and disseminates circRNA-related sequence data and fundamental details, ensuring public availability (34). The cellular localization of circRNAs was obtained from the RNALocate database, which compiles comprehensive RNA-seq data across various tissues and RNA types, including circRNAs (35).

#### 3.2.1. ALS gene expression pattern

To have an insight into gene expression in different tissues in ALS, we used the GSE153960 dataset from the GEO database (https://www.ncbi.nlm.nih.gov/geo/). This dataset contains 1,838 samples from cerebellum (n=331), cortex motor lateral (n=137), cortex motor medial (n=137), cortex motor unspecified (n=74), cortex occipital (n=96), cortex sensory (n=2), cortex temporal (n=102), frontal cortex (n=327), hippocampus (n=53), spinal cord cervical (n=258), spinal cord lumbar (n=239), and spinal cord thoracic (n=82) of ALS patients and non-neurological (control). In this study, all the samples were used other the cortex sensory and cortex motor unspecified. Then, the data was analyzed by GEO2R, the online tool on the GEO database. A threshold of FC ≥ 1 and p-value < 0.05 was considered for the result of RNA-seq data.

#### 3.2.2. circRNAs-miRNA and miRNA-mRNA interaction prediction

To achieve this, the circAtlas 3.0 database (https://ngdc.cncb.ac.cn/circatlas/), a robust platform for circRNA analysis, was utilized. Accordingly, circRNA sequences were entered into circAtlas to forecast potential interactions with miRNAs. CircAtlas integrates three distinct prediction tools—TargetScan, miRanda, and PITA—to identify possible circRNA-miRNA interactions. In this study, only miRNAs predicted by at least two of these tools were selected for further examination. (36). The association between the identified miRNAs and diseases was validated using HMDD 4 (https://www.cuilab.cn/hmdd), a database that compiles experimentally confirmed human miRNA-disease relationships. Consequently, miRNAs linked to ALS were extracted from HMDD 4 and subsequently compared with those targeted by ALS-associated circRNAs (37). To explore possible gene targets of miRNAs and AD-related miRNAs influenced by AD-associated circRNAs, we utilized the MIENTURNET database (http://userver.bio.uniroma1.it/apps/mienturnet/). In our analysis, only results with a p-value below 0.05 were deemed statistically significant (38). Furthermore, to investigate possible links between the target genes of miRNAs and human health conditions, the DisGeNET database (https://www.disgenet.com/) was utilized. In this approach, ALS-related genes (ID: C0002736) were retrieved from DisGeNET and analyzed alongside miRNAs’ target genes for comparison (39).

#### 3.2.3. Enrichment analysis

RummaGEO is an online platform that enables researchers to explore gene expression patterns across a comprehensive set of RNA sequencing (RNA-seq) studies available in the GEO database. Covering both human and mouse datasets, RummaGEO (https://rummageo.com/) was employed to analyze common gene targets of miRNAs. Then, RummaGEO was employed to identify biological processes and the specific cell types influenced by common gene targets of miRNAs that are sponged by AD-related circRNAs. (40). To investigate the cellular functions of miRNA target genes and circRNA-associated protein targets, pathway enrichment analysis was performed. The Enrichr tool (https://maayanlab.cloud/Enrichr/) was utilized to identify relevant biological pathways by comparing miRNA target genes and circRNA protein interactions with established KEGG pathways. Only pathways with a p-value below 0.05 were deemed statistically significant (41).

#### 3.2.4. ORF analysis

To find the possibility of translation of ALS-related circRNAs to proteins, the circAtlas database was employed. In this way, the sequence of the circRNAs was uploaded into circAtlas, and then the sequence of proteins which can be generated from these circRNAs was extracted. In the next step, the sequence of circRNAs was blasted in the Uniprot BLAST online tool (https://www.uniprot.org/blast). Only the proteins with 100% similarity with the peptides’ sequence were selected.

#### 3.2.5. protein target prediction for circRNAs

CircNetVis (https://www.meb.ki.se/shiny/truvu/CircNetVis/) was utilized to examine potential interactions between circRNAs and proteins. To achieve this, circRNA sequences in FASTA format were submitted to CircNetVis, and the resulting protein interaction data were retrieved for further analysis (42). Subsequently, pathway enrichment analysis was conducted using the Enrichr database, while the STRING database was employed to construct the protein-protein interaction (PPI) network (43).

#### 3.2.6. Molecular Docking

Since primary data showed hsa_circ_0060762, hsa_circ_0023919, hsa_circ_0000567, hsa_circ_0008870, hsa_circ_0004846, hsa_circ_0034880, and hsa_circ_0005171 can interact with FUS, and this protein is important in ALS development, the interaction of FUS and the circRNAs was verified by molecular docking. However, the length of these circRNAs is more than 500nt, which is higher than the limitation of most bioinformatic tools. Therefore, we extracted the most important part of each circRNA with FUS. In this manner, RBPsuit 2.0 was used for finding the important area of the circRNAs in interaction with FUS. This web server divides each sequence into 101nt and verifies its interaction with RBPs. In addition, it supports circRNA sequences in different organisms, as well as human (44). Then, each segment that had at least a 0.6 score was selected for molecular docking. In the next step, the crystal structure of the wild type of FUS (PDB ID: 5YVI) and mutant (P525L) FUS (PDB ID: 7CYL) was downloaded in PDB format from the Protein Data Bank (https://www.rcsb.org/) (45). The crystal structure was set up for docking in Chimera 1.8.1 by fixing incomplete residues, adding hydrogen atoms, and removing water and ligands (46). Finally, the interaction of FUS and selected segments was verified by the HDOCK web server (47).

## 4. Results

Through a search of the specified keywords in PubMed, Web of Science, and Scopus, 140 records were initially identified. After eliminating duplicates, the number was reduced to 84. The titles of the articles were reviewed, resulting in the exclusion of 38 studies that were not related to MND, such as those focusing on cancer, Alzheimer’s disease, and other unrelated conditions. In the screening phase, 29 review papers were excluded, followed by the verification of data from full-text articles. Just one preprint article was found and removed. In the final stage, 14 articles were deemed eligible, with 28 circRNAs associated with ALS and 3 circRNAs with SMA being identified.

**Table 1.**
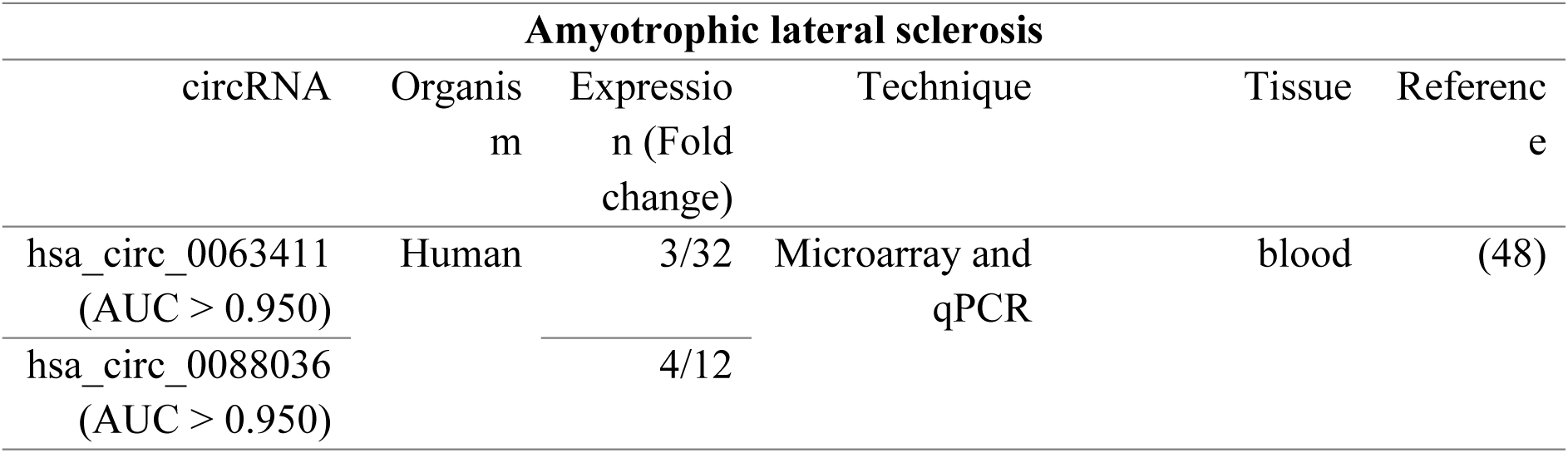
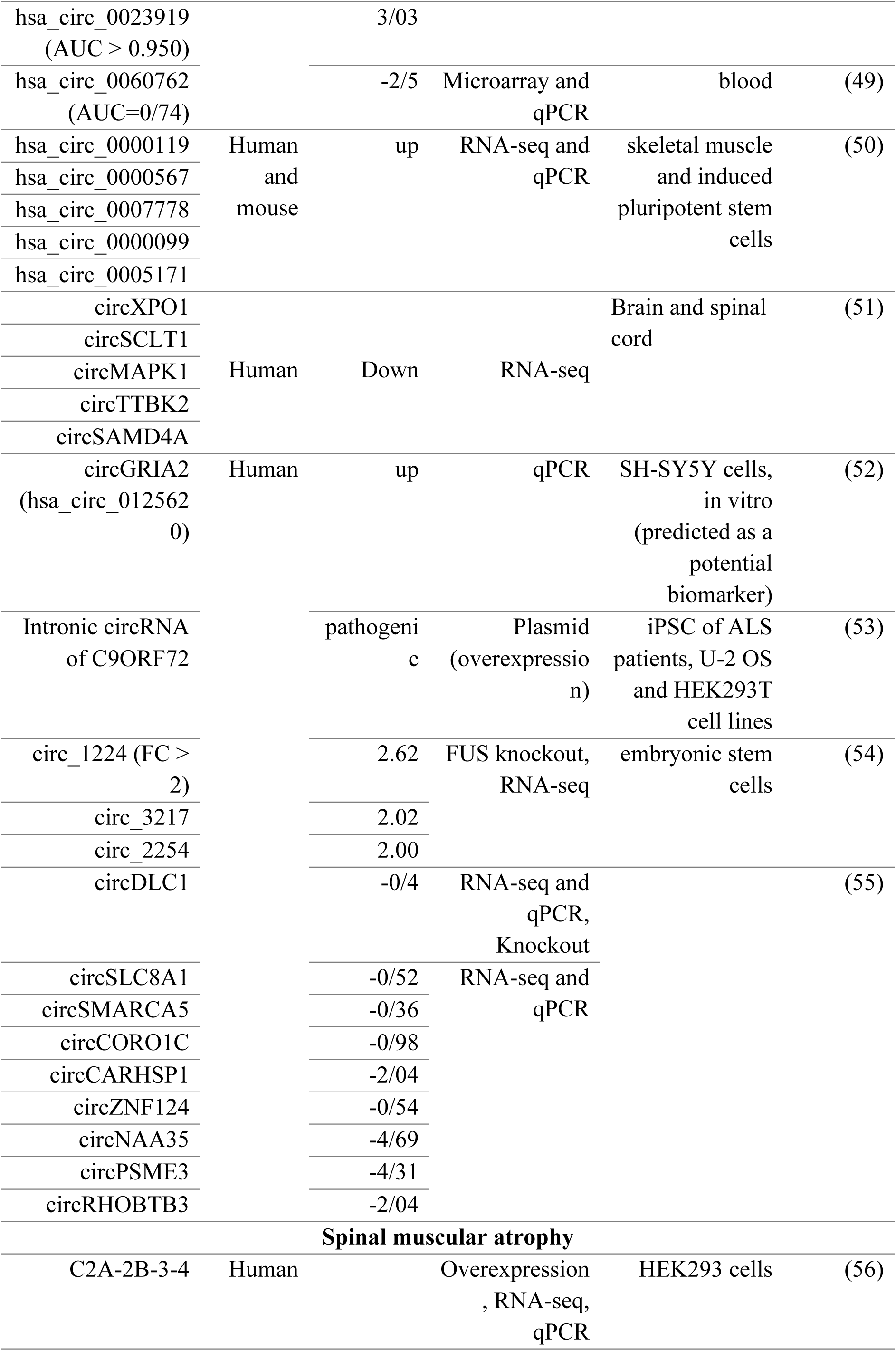
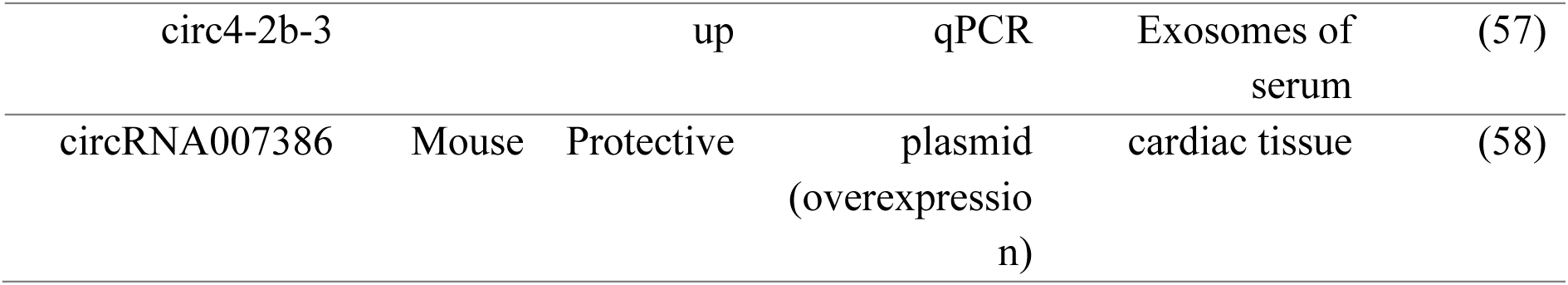
The list of circRNAs that was extracted from the literature review.

### 4.1. Amyotrophic Lateral Sclerosis CircRNAs

#### 4.1.1. circRNAs expression

A study on three tissue types: spinal cord cervical, spinal cord thoracic, and spinal cord lumbar of ALS patients has revealed that the expression of a high number of circRNAs significantly these three tissues in ALS. These numbers of circRNAs were 11,702, 9976, and 8481 for spinal cord cervical, spinal cord thoracic, and spinal cord lumbar, respectively (59). Another study on mice and brain tissue samples from post-modern ALS patients has identified 30 circRNAs related to ALS in mice and 18 circRNAs in the brain of ALS patients. Among the 18 circRNAs, 9 of them significantly downregulate, and none of them significantly upregulate and five of them mostly downregulate, including circXPO1, circSCLT1, circMAPK, circTTBK2, and circSAMD4A (51).

#### 4.1.2. Biomarkers

Due to the stability of circRNAs, they can be a good choice as a biomarker of ALS. While two circRNAs, including hsa_circ_0063411 and hsa_circ_0088036 upregulated, hsa_circ_0023919 is downregulated in the blood of ALS patients. The three circRNAs demonstrated a high specificity and sensitivity, highlighting their strong potential as diagnostic biomarkers (48). Another study conducted on both male and female ALS patients identified hsa_circ_0060762 as a potential biomarker for ALS in the patients’ blood, which originates from the Chromosome Segregation 1 Like (CSE1L) gene (49).

In ALS, RNA-sequencing and RT-qPCR revealed changes in circRNA levels in skeletal muscle and motor neurons, with notable differences across disease stages and tissues. Some circRNAs elevated in ALS muscle were reduced in spinal cord samples, including hsa_circ_0000119, hsa_circ_0000567, hsa_circ_0005171, hsa_circ_0047886, hsa_circ_0001948, and hsa_circ_0000033, offering insights into neuromuscular molecular programs and therapeutic strategies (50).

In sporadic ALS, reduced RNA editing at the GluA2 mRNA site is linked to ADAR2 downregulation, which plays a critical role in disease pathogenesis. TDP-43 pathology is observed in motor neurons lacking ADAR2 in both ALS patients and ADAR2 knockout mice. Extracellular RNAs with ADAR2-dependent editing sites, found in body fluids, may serve as potential ALS biomarkers. Research identified 10 such sites in RNAs from SH-SY5Y cells and their medium, including the arginine/glycine site of SON mRNA and a circRNA, including circGRIA2 (hsa_circ_0125620) with an ADAR2-dependent site. Changes in editing efficiency at these sites might indicate ALS biomarker potential (52).

#### 4.1.2. circRNA generated from the C9ORF72 gene increases toxicity in motor neurons

C9ORF72 hexanucleotide repeat expansion is the most frequent genetic factor behind ALS and FTD. Toxicity is mediated by repeat-containing RNA through nuclear granules and dipeptide repeat proteins. Using single-molecule imaging, researchers discovered that G-rich repeats form intronic circRNA due to defective lariat processing, which serves as the translation template. The Nuclear RNA Export Factor 1 (NXF1)-Nuclear Transport Factor 2 Like Export Factor 1 (NXT1) pathway facilitates the export of this circular RNA to the cytoplasm and influences dipeptide repeat production and accumulation (53).

#### 4.1.3. FUS regulates expression of circRNAs

The RNA-binding protein Fused in sarcoma (FUS) is involved in various RNA biosynthesis processes and is associated with the development of ALS and frontotemporal dementia. Through RNAi and the overexpression of wild-type and ALS-linked FUS mutants, it has been demonstrated that circRNA biogenesis can be influenced by changes in FUS nuclear levels and potential toxic gain-of-function effects. FUS controls circRNA biogenesis by interacting with introns adjacent to back-splicing junctions, and this regulation can be replicated using artificial constructs (54).

The impact of the P525L FUS mutation on unconventional splicing processes that result in the generation of circRNAs. The P525L FUS mutation in motor neurons leads to disruptions in circRNA expression by binding to introns adjacent to downregulated circRNAs, particularly those containing inverted Alu repeats. Nine circRNAs including circDLC1 (p-value just above the significance level), circSLC8A1, circSMARCA5, circCORO1C, circCARHSP1, circZNF124, circNAA35, circPSME3, and circRHOBTB3 have reported downregulate in motor neurons of ALS patients. Except circCARHSP1 and circPSME3, the other seven circRNAs upregulate during motor neuron differentiation. In this context, circDLC1 is downregulated by the P525L FUS mutation in motor neurons, despite being one of the most abundant circRNAs in human motor neurons. Experimental evidence suggests that the downregulation of circDLC1 leads to a reduction in ENAH Actin Regulator (ENAH) level. In addition, The P525L FUS mutation leads to a significant delocalization of circRNAs within the cytoplasm of motor neurons, including circCARHSP1, circNAA35, and circPSME3 are shifted towards the cytoplasm from the nucleus in FUSP 525L (55).

### 4.2. Spinal muscular atrophy

#### 4.2.1. C2A-2B-3-4 is a potential biomarker of SMA

The SMN1 gene encodes SMN, a vital protein for RNA metabolism, and its loss due to deletions or mutations causes SMA. While SMN2, a similar gene, cannot fully compensate due to exon 7 skipping, therapies targeting splicing or gene replacement restore SMN. SMN genes produce numerous circRNAs. Experiments in HeLa and HEK 293 cells revealed that intronic sequences and forward splicing enhance the creation of circRNAs containing multiple exons, often localized in the cytoplasm (60).

SMN1 and SMN2 (collectively SMN1/2) produce circular RNAs, including the abundantly expressed C2A-2B-3-4, which spans exons 2A, 2B, 3, and 4. Overexpression of C2A-2B-3-4 in HEK293 cells altered the expression of ∼15% of genes and affected 61 proteins. These changes differed significantly from those caused by L2A-2B-3-4, a linear transcript. C2A-2B-3-4 influenced genes related to chromatin remodeling, transcription, lipid metabolism, and neuromuscular junctions. This highlights its distinct regulatory role and broad impact on SMN1/2 gene functions (56). SMN circ4-2b-3 is the only SMN circRNA detected in both type I SMA cell exosomes and patient serum. A subgroup of type I SMA patients, called super-responders, shows high levels of SMN circ4-2b-3 and responds well to Nusinersen therapy. Nusinersen is an intrathecal antisense oligonucleotide that corrects the SMN2 splicing defect.

Therefore, SMN circ4-2b-3 serves as a potential biomarker to predict type I SMA patients’ response to Nusinersen (57). SMN is one of the human genes with the highest density of Alus, which are evolutionarily conserved in primates and frequently appear in an inverted orientation. Inverted repeat Alus (IRAlus) hinder the splicing of long introns in SMN while facilitating the extensive formation of alternative circRNAs. The protein Sam68 attaches near Alus in the SMN pre-mRNAs and enhances the formation of circRNAs (61).

#### 4.2.2. circRNA007386, as a protective circRNA in SMA

Transcriptome sequencing in a severe SMA mouse model revealed thousands of differentially expressed RNAs, including miR-34a deregulation across tissues. An in vitro study has revealed that reducing SMN expression led to an increase in miR-34a-5p levels, while elevated miR-34a-5p independently interfered with cell-cycle progression by modulating its targets, mirroring the gene expression profiles seen in SMA mouse cardiac tissue. Moreover, an inhibitor of this miRNA increases circRNA007386 expression, which originates from the RyR2 gene. However, circRNA007386 contains two target sites for sponging miR-34a-5p. miR-34a-5p emerges as a key contributor to cardiac pathologies in the SMA mouse model, alongside circRNA007386 and three other elements, including Birc5, Spag5, and possibly lncRNA00138536 (58).

### 4.3. In silico analysis

We conducted a bioinformatics analysis of human circRNAs associated with MND, each identified by a circBase ID (Table 2). Therefore, bioinformatic analysis was conducted for 15 circRNAs in ALS, including hsa_circ_0063411, hsa_circ_0088036, hsa_circ_0023919, hsa_circ_0060762, hsa_circ_0000119, hsa_circ_0000567, hsa_circ_0007778, hsa_circ_0000099, hsa_circ_0005171, hsa_circ_0125620, hsa_circ_0001017, hsa_circ_0001439, hsa_circ_0008870, hsa_circ_0034880, and hsa_circ_0004846. Among them, hsa_circ_0000099 had the longest and hsa_circ_0088036 the shortest lengths by 1307bp and 279, respectively. According to the circBase database, all 15 circRNAs are expressed in the nervous system and related cells, other than hsa_circ_0125620 (Table 2). Almost all of them are expressed in extracellular vesicles. However, no tissue expression and subcellular localization were found for hsa_circ_0125620.

**Table 2.**
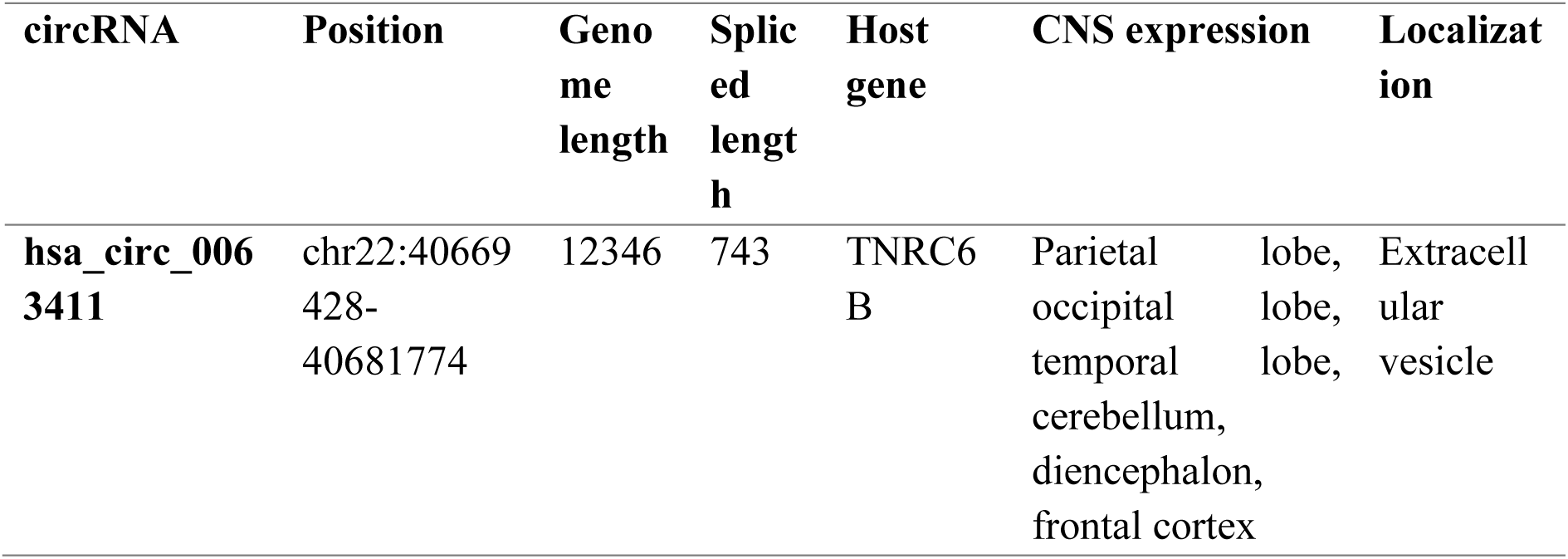
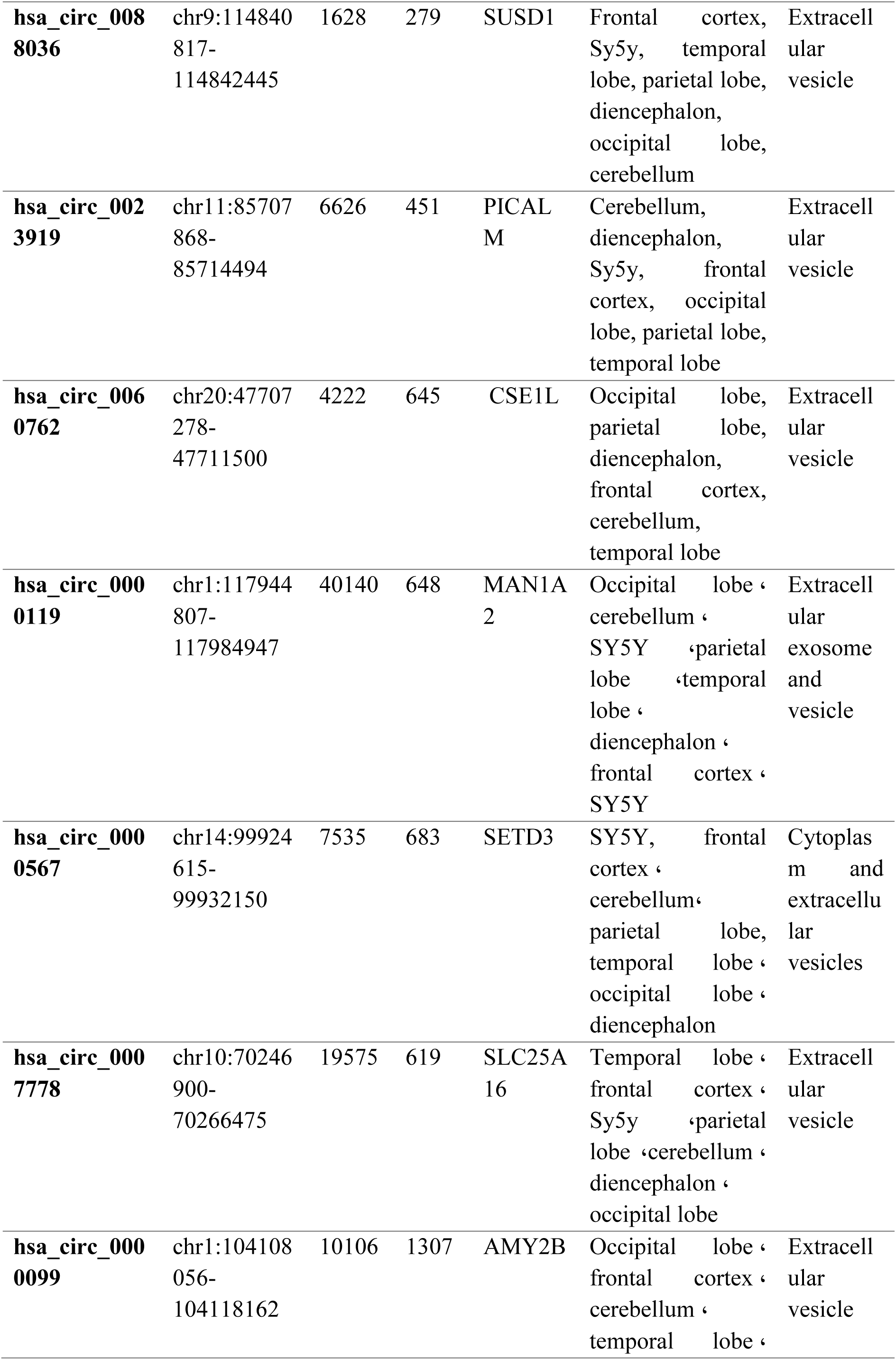
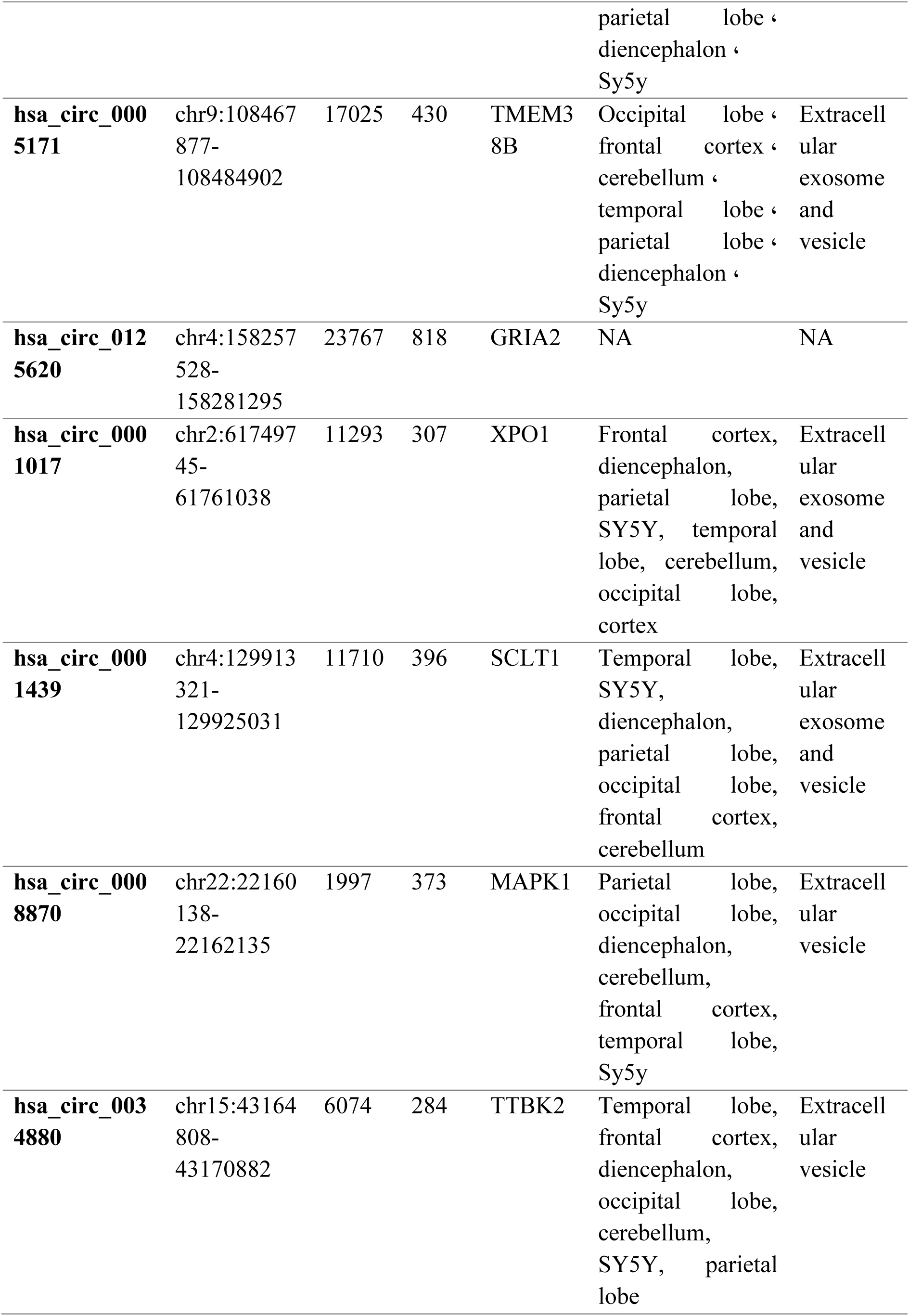
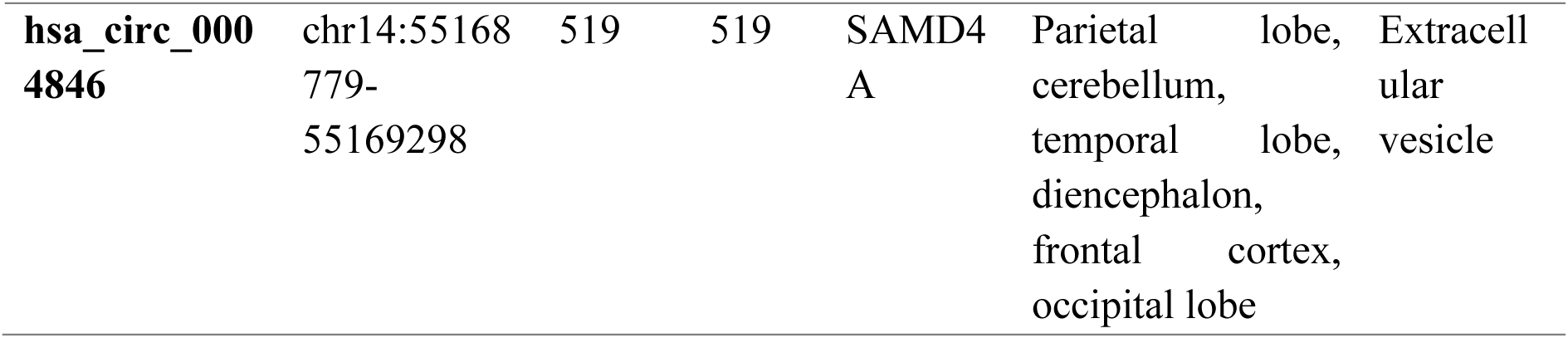
ALS-associated circRNAs. This table gives basic information about ALS-related circRNAs, including genomic position and length, their length after splicing, tissue expression, and subcellular localization.

#### 4.3.1. Pathway enrichment analysis for targets of selected circRNAs

The miRNA targets of ALS-related circRNAs are predicted by circAtlas 3.0. However, none of them were found as ALS-related miRNA, based on the HMDD 4.0 database. In the next step, the gene target of miRNAs that sponge with ALS-related circRNAs was predicted by MIENTURNET.

Then, the relation of these genes with ALS was verified by the DisGeNET database (Figure 3A). Among the 10 circRNAs, hsa_circ_0000099, hsa_circ_0001017, hsa_circ_0004846, and hsa_circ_0034880 had the highest number of ALS-associated genes. Pathway enrichment analysis was also performed for all 15 circRNAs (Figure 3B). hsa_circ_0000099 showed a high number of genes in neuroactive ligand-receptor interaction, cytokine-cytokine receptor interaction, and calcium signaling pathway. Long-term potentiation was a common cellular pathway for hsa-_circ_0000099, hsa_circ_0000567, hsa_circ_0060762, hsa_circ_0063411, and has_circ_0034880. Wnt signaling pathway was also common among hsa_circ_0005171, hsa_circ_0060762, hsa_circ_0063411, and hsa_circ_0001017. hsa_circ_0000099, hsa_circ_0005171, hsa_circ_0023919, and hsa_circ_0008870 target PPAR signaling pathway. Wnt signaling pathway was also common among hsa_circ_0023919, hsa_circ_0060762, hsa_circ_0063411, and hsa_circ_000870.

**Figure 3.**
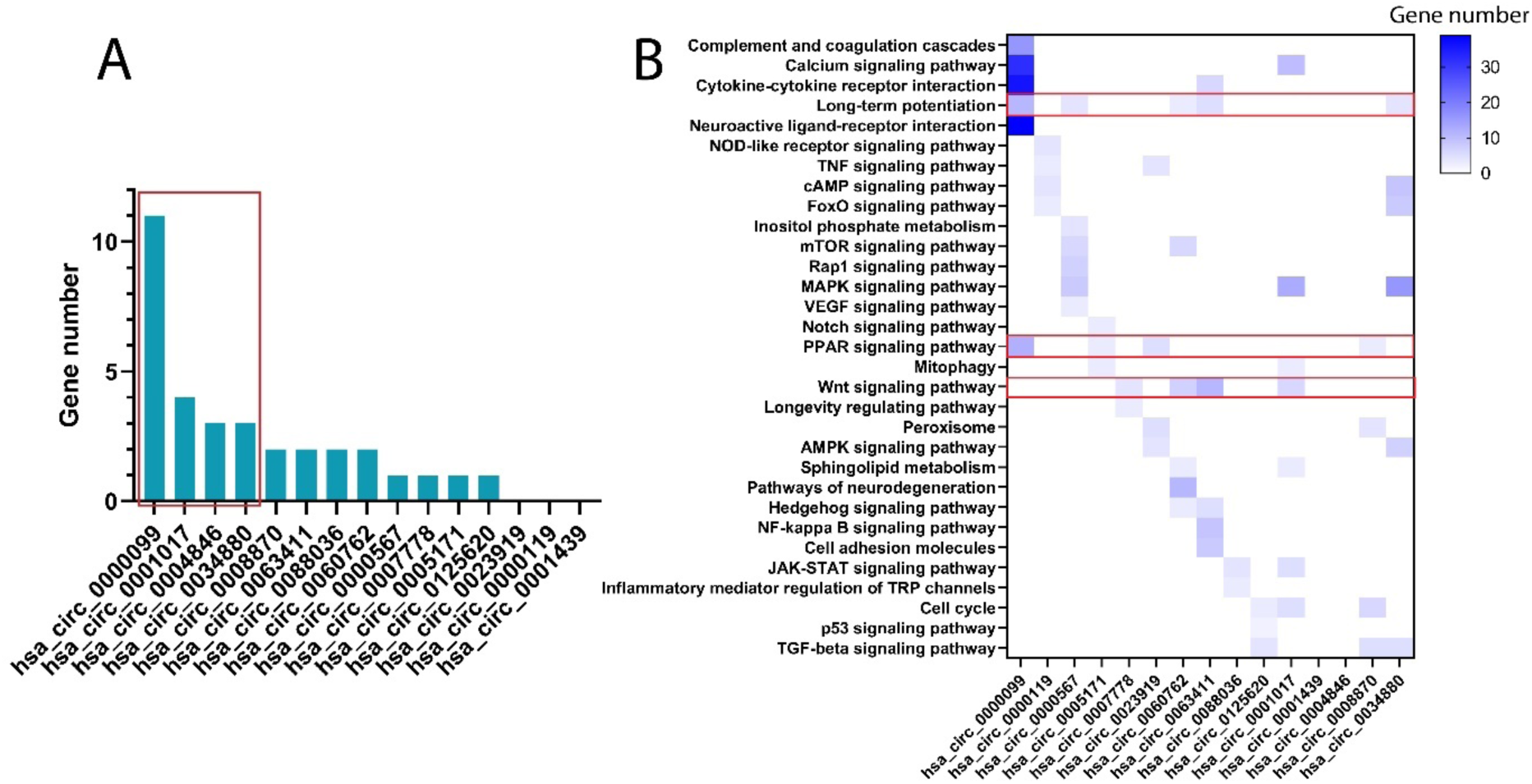
Analysis of ALS-related circRNAs. **A)** A total of 11 and 4 ALS-related genes were found for miRNAs which target by hsa_circ_0000099 and hsa_circ_0001017. This figure was 4 for has_circ_004846 and has_circ_0034880. **B)** cellular pathways for the gene target of miRNAs which sponged by ALS-related circRNAs. Long-term potentiation, PPAR, and Wnt signaling pathways were associated with a high number of circRNAs.

#### 4.3.2. Analysis of circRNAs common gene targets

To clarify the distinct roles of specific circRNAs in ALS, we analyzed the shared gene targets of miRNAs linked to each circRNA (Figure 4A). In this manner, three highest raked genes were selected for each circRNAs. Eight target genes including MDM2 proto-oncogene (MDM2), CREB3 regulatory factor (CREBRF), TATA-box binding protein associated factor 8 (TAF8), solute carrier family 35, member E2 (SLC35E2), trans-golgi network vesicle protein 23 homolog C (TVP23C), serine/threonine kinase 4 (STK4), MAPK activated protein kinase 5 (MAPKAPK5), and stearoyl-CoA desaturase (SCD) were found for hsa_circ_0000099. NPL4 homolog, ubiquitin recognition factor (NPLOC4), transmembrane protein 184B (TMEM184B), aldolase, fructose-bisphosphate A (ALDOA), collagen and calcium binding EGF domains 1 (CCBE1), tyrosine 3-monooxygenase/tryptophan 5-monooxygenase activation protein epsilon (YWHAE), midnolin (MIDN), trinucleotide repeat containing adaptor 6B (TNRC6B), and solute carrier family 7 member 5 (SLC7A5) were common gene targets for hsa_circ_0004846. LIF interleukin 6 family cytokine (LIF), pantothenate kinase 1 (PANK1), solute carrier family 43 member 2 (SLC43A2), glycerol kinase 5 (GK5), ankyrin repeat and SOCS box containing 16 (ASB16), UBX domain protein 2A (UBXN2A), and gigaxonin (GAN) were also obtained as the common gene targets for hsa_circ_0034880. However, no common gene target was found for has_circ_0001017.

**Figure 4.**
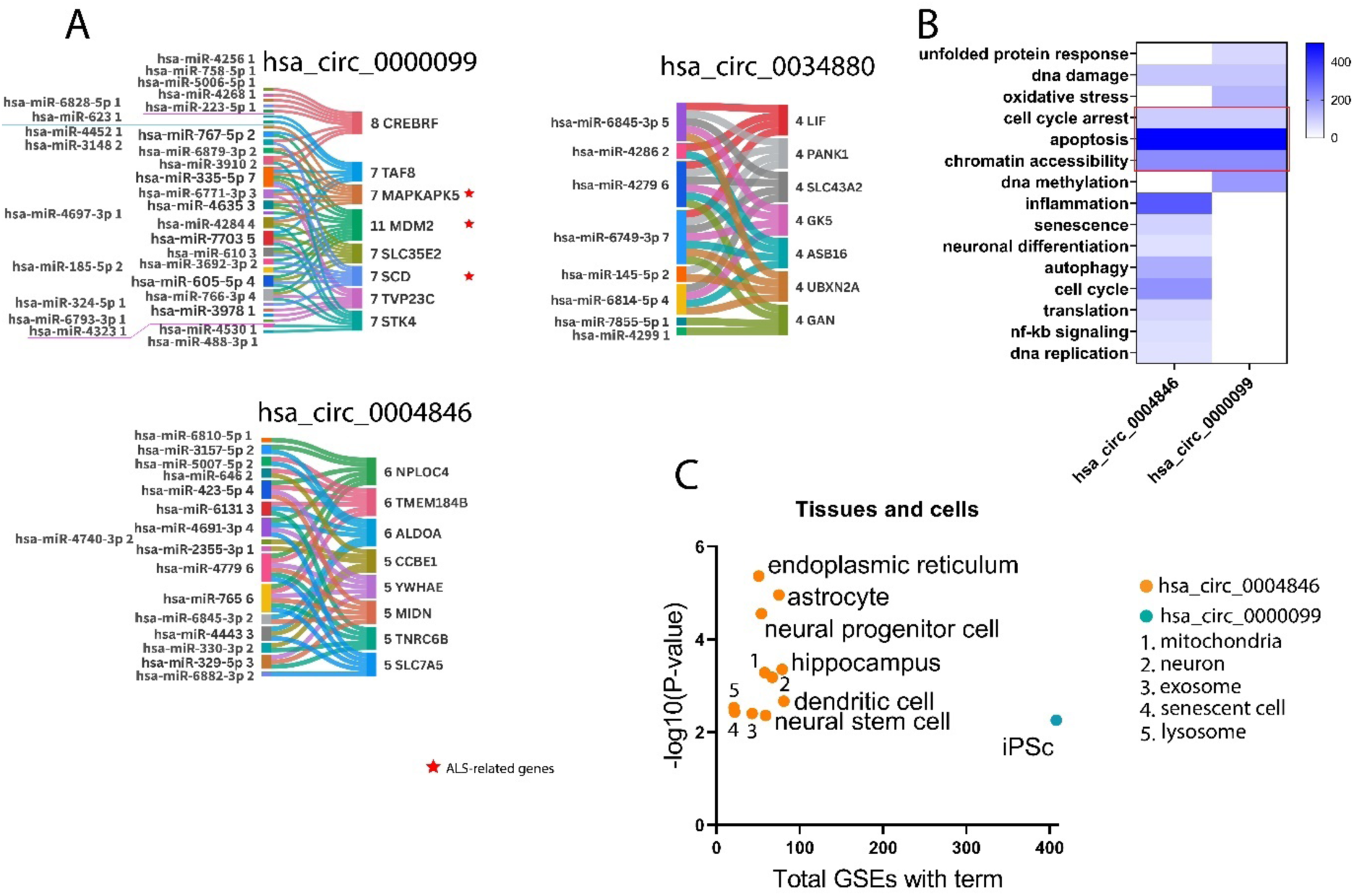
Analysis of common gene targets of AD-related miRNAs sponged by hsa_circ_0000099, hsa_circ_0004846, and hsa_circ_0034880. **A)** The common gene target of ALS-related circRNAs. **B)** The cellular pathways/biological process of the common gene targets. Cell arrest, apoptosis, and chromatin accessibility were common between hsa_circ_0000099 and hsa_circ_0004846. **C)** RummaGEO analysis for brain cells revealed that hsa_circ_0004846 is potentially involved in neuronal and astrocytes functions.

Gene enrichment analysis was performed for common gene targets of hsa_circ_0000099, hsa_circ_0034880, and hsa_circ_0004846 (Figure 4B). According to the results, biological process/cellular pathways of apoptosis, cell cycle arrest and chromatin accessibility were common between hsa_circ_0000099 and hsa_circ_0004846. Expression analysis also was performed for the common genet targets to verified these genes expression pattern in which related cells (Figure 4C). Several cells and tissues were obtained for has_circ_0004846, including endoplasmic reticulum, astrocytes, neuronal progenitor cell, hippocampus, mitochondria and neuron, while just iPSc was found for has_circ_0000099.

#### 4.3.3. ORF analysis

To verify whether the ALS-related circRNAs could be translated into protein, ORF analysis was performed for all 15 ALS-related circRNAs. According to the results, none of them were able to translate to a full-length protein. However, the longest peptide can be translated from has_circ_0000099 by 367aa, which is exactly similar to the first 367 amino acids of Alpha-amylase 2B (AMY2B) (Figure 5).

**Figure 5.**
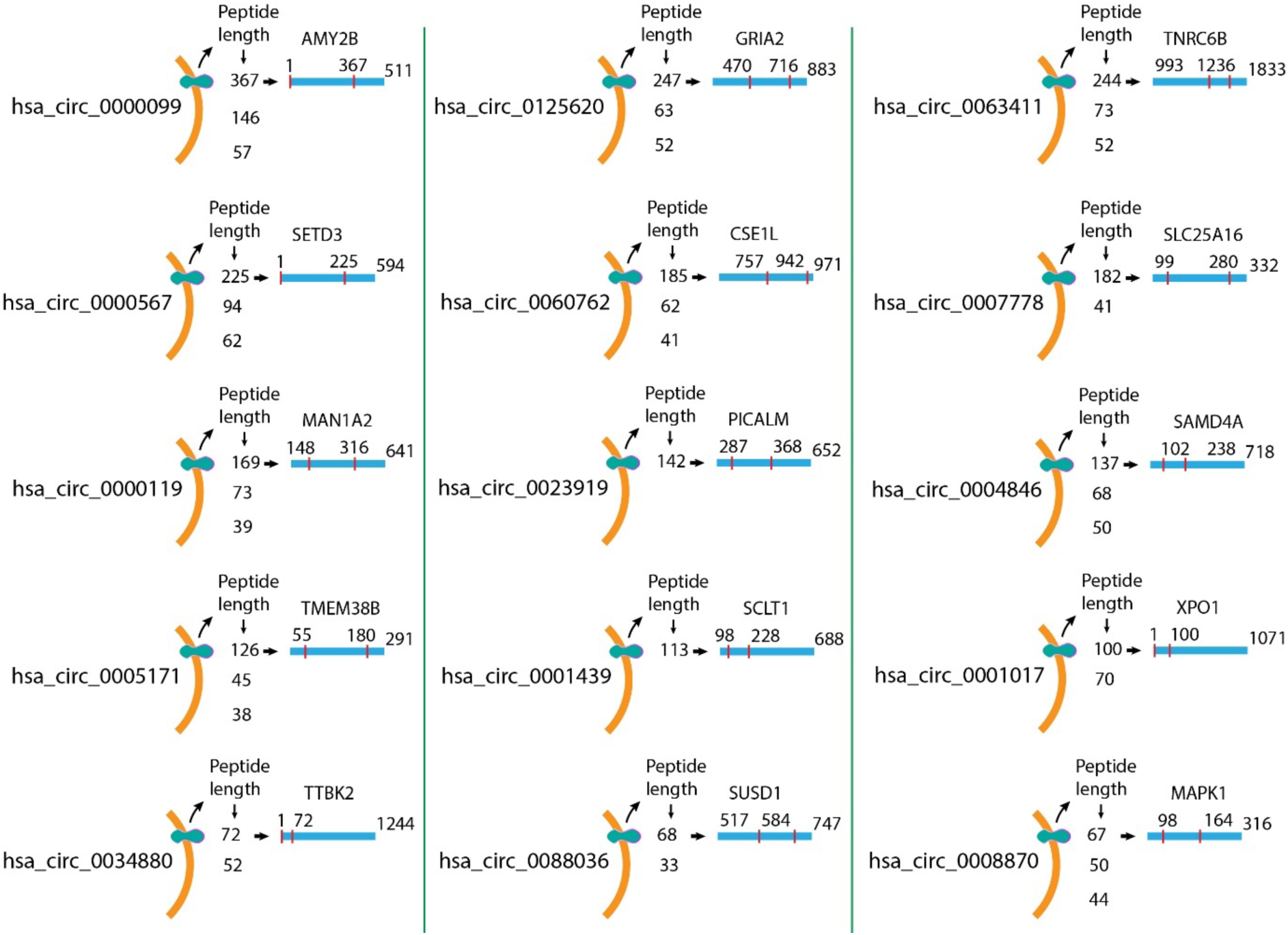
ORF analysis for the ALS-related circRNAs. According to the data, no full-length protein is generated from these circRNAs.

#### 4.3.4. circRNAs-protein interactions

The interaction of ALS-related circRNAs and proteins was predicted by circNetVis (Figure 7). Among the 10 circRNAs, hsa_circ_0060762 exhibited the highest number of interactions, engaging with 15 proteins including eukaryotic initiation factor 4A-III (EIF4A3), Protein argonaute-2 (AGO2), insulin-like growth factor 2 mRNA-binding protein 2 (IGF2BP2), FMR1-interacting protein NUFIP2 (FMRP), polypyrimidine tract-binding protein 1 (PTB), ELAV-like protein 1 (ELAV1), insulin-like growth factor 2 mRNA-binding protein 3 (IGF2BP3), insulin-like growth factor 2 mRNA-binding protein 1 (IGF2BP1), microprocessor complex subunit DGCR8 (DGCR8), Nucleolysin TIAR (TIAL1), protein lin-28 homolog A (LIN28A), RNA-binding protein FUS (FUS), protein argonaute-1 (AGO1), Zinc finger CCCH domain-containing protein 7B (ZC3H7B), splicing factor U2AF 65 kDa subunit (U2AF65) (Figure 7A). Hsa_circ_0000119 and hsa_circ_0008870 interact with 13 and 11 proteins, respectively. Moreover, hsa_circ_0034880, hsa_circ_004846, hsa_circ_0023919, hsa_circ_0005171, hsa_circ_008870, hsa_circ_0000567, and hsa_circ_0060762 can interact with FUS protein (Figure 6A). The interactions of proteins with each other were verified by STRING database and analyzed by Cytoscape. Results show that FMR1, ELAV1, and IGF2BP1 are the hub proteins in this protein-protein interaction network (Figure 6B). FMR1 can interact with hsa_circ_0000119, hsa_circ_0023919, hsa_circ_0060762, and hsa_circ_0063411 ALS-related circRNAs (Figure 7A). Pathway enrichment analysis was performed for all proteins that can interact with ALS-related circRNAs (Figure 6C). Three cellular pathways, including mRNA surveillance, spliceosome, and RNA transport cellular pathways, were found for these proteins.

**Figure 6.**
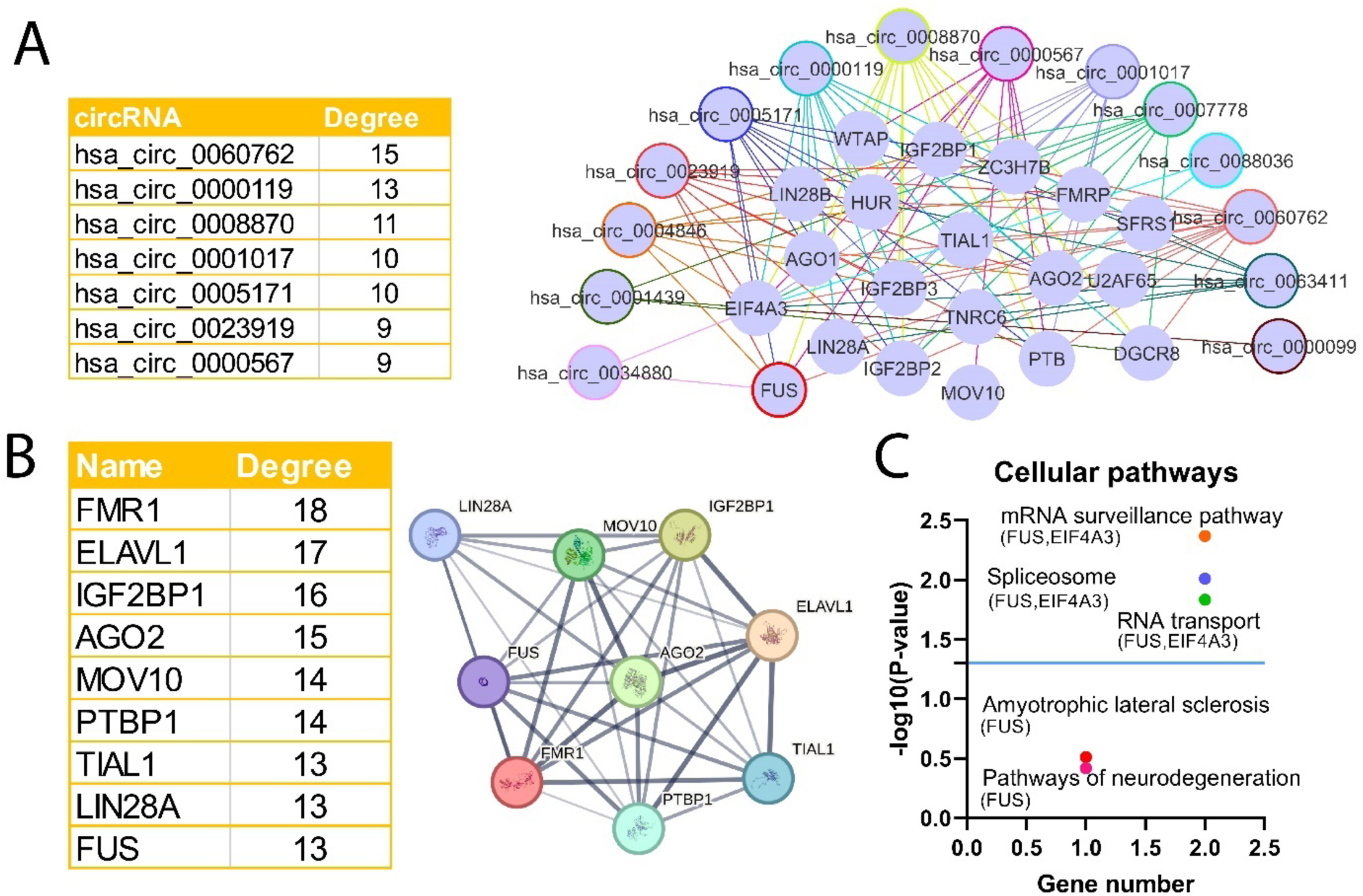
Interaction of ALS-related circRNAs and proteins. **A)** hsa_circ_0060762, hsa_circ_0000119, and hsa_circ_0008870 had the highest interaction with proteins by 15, 13, and 11 interactions, respectively. **B)** PPI network for proteins with interact with ALS-related circRNAs. FMR1, ELAVL1, IGF2BP1, and AGO2 were the hub proteins.

**Figure 7.**
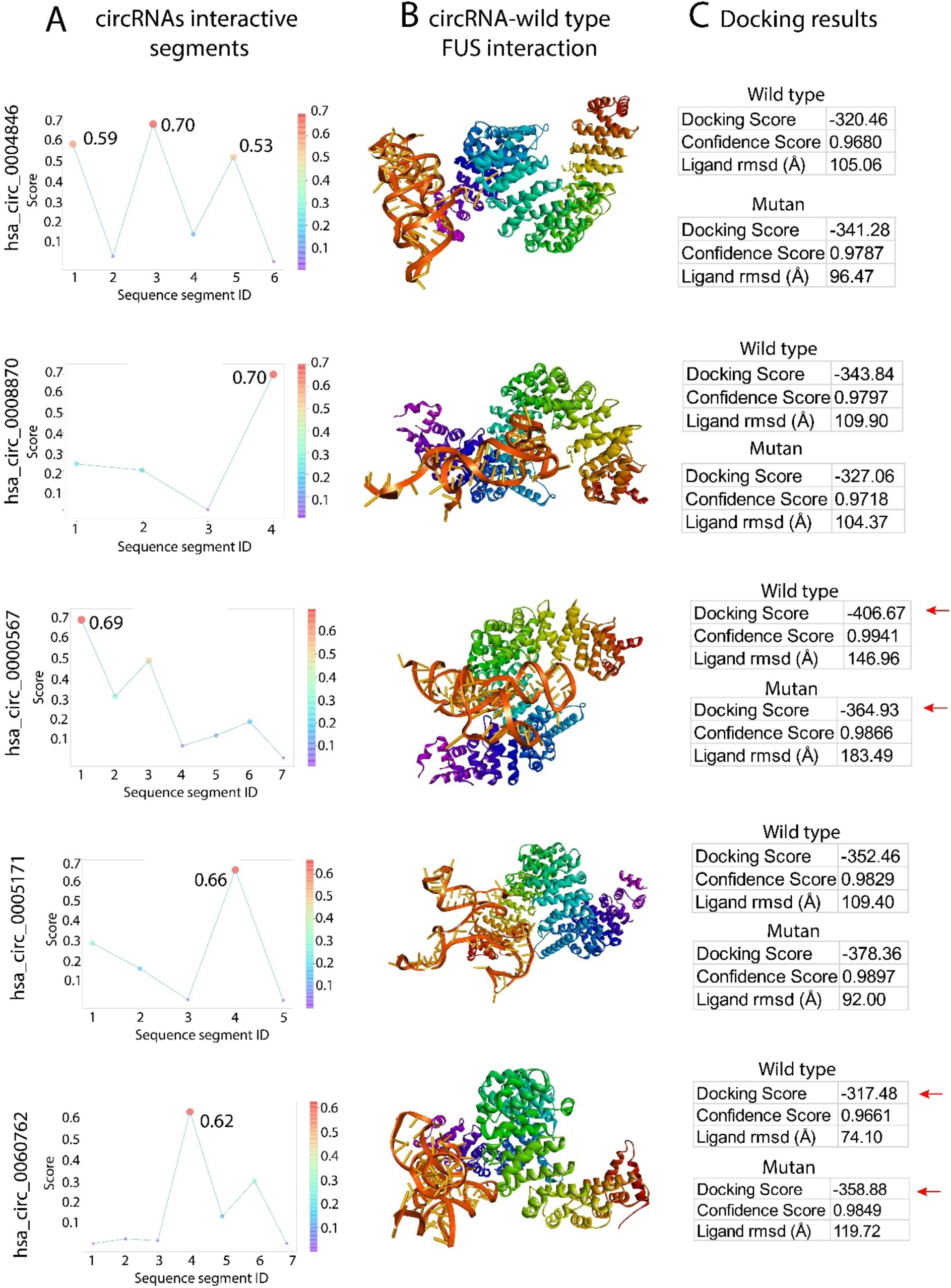
Molecular docking for wild-type and mutant FUS with hsa_circ_004846, hsa_circ_0008870, hsa_circ_0000567, hsa_circ_0005171, and hsa_circ_0060762. **A)** The line plot shows the interaction score of each segment of circRNAs with FUS protein. **B)** Interaction of circRNAs with wild-type FUS. **C)** The result of docking for the interaction of selected segments and the FUS protein. Below each table is the sequence of selected segments. Mutation of FUS noticeably changes its affinity to hsa_circ_0000567 and hsa_circ_0060762.

### 4.4. Molecular docking

The result of circNetVis showed that hsa_circ_0060762, hsa_circ_0023919, hsa_circ_0000567, and hsa_circ_0005171 can interact with the FUS protein. Due to the importance of FUS protein in the development of ALS, interactions of these circRNAs with wild-type and mutant FUS (P525L) were verified by molecular docking (Figure 7). According to the results of RBPsuite, three parts of hsa_circ_0004846 can interact with wild-type FUS, and the bending score of the third segment was 0.70 (Figure 7A). hsa_circ_0008870 can also interact with wild-type FUS with its segment 4, with a score of 0.70. This value was 0.69 for the first segment of hsa_circ_0000567. Both hsa_circ_0005171 and hsa_circ_0060762 can interact with wild-type FUS with their fourth segment, with binding scores of 0.66 and 0.62, respectively. Molecular docking was performed on ALS-related circRNAs that exhibited a one segment binding score of at least 0.6 in their interaction with FUS. In this step, these segments were selected to molecular docking analysis to verified the interaction of these circRNAs with wild-type and mutant FUS (P525L) (Figure 7B). Due to molecular docking results, hsa_circ_0004846 interacts with wild-type FUS with a −320.46 docking score. However, this affinity increases by −21 docking score with mutant FUS (Figure 7C). This amount was −343.84 and −327.06 docking score for the interaction of hsa_circ_0008870 with wild-type and mutant FUS, respectively. A mutation in FUS can significantly change hsa_circ_0000567 affinity to this protein. According to the molecular docking results, the affinity of hsa_circ_0000567 with wild-type FUS is −406.67 docking score, which is 41.74 stronger than that of mutant FUS. The docking score of the hsa_circ_0005171 interaction with wild-type FUS was −352.46, while it was −378.36 with mutant FUS. The changing affinity of hsa_circ_0060762 with wild-type FUS noticeably changed with a mutation in FUS. hsa_circ_0060762 interacts with wild-type FUS with a −317.48 docking score. The corresponding value was −358.88 for the interaction of hsa_circ_0060762 with the mutant FUS.

### 4.5. RNA-seq data analysis

In order to obtain the gene expression pattern of ALS in different tissues, we used RNA-seq data on the GEO database. The result of gene expression was different for each tissue. The highest gene expression change was seen in the cervical spinal cord and the lowest in the temporal cortex, respectively. Since the rate of gene expression change was not significant in the temporal and frontal cortex, they were excluded from further analysis.

Based on FC, the top 10 genes are illustrated in each tissue (Figure 8A). Albumin (ALB), tRNA-Lys (anticodon CTT) 2-2 (TRK-CTT2-2), LOC107985634, guanylate cyclase activator 1C (GUCA1C), and C-C motif chemokine ligand 18 (CCL18) exhibited the highest expression in the cerebellum, lateral motor cortex, medial motor cortex, hippocampus, and cervical spinal cord, respectively. Additionally, LOC107985634, CCL18, and chitinase 1 (CHIT1) showed the highest expression in the occipital cortex, lumbar spinal cord, and thoracic spinal cord, respectively. Several genes appear in multiple regions within the dataset. Notably, LOC107985853 is present in the medial motor cortex, occipital cortex, and cervical spinal cord. Additionally, genes such as CHIT1, CCL18, LINCO132, and LYZ are shared across multiple regions. Pathway enrichment analysis was also performed for upregulated genes (Figure 8B). However, no ALS-related cellular pathways were obtained for the spinal cord, occipital, and cortex motor medial. Hippocampus showed a wide range of neurodegenerative-related cellular pathways, including pathways of neurodegeneration, ALS, Alzheimer’s, and Parkinson’s disease. Neuroactive ligand-receptor interaction was common among the hippocampus, cervical, and lumbar spinal cords. cell adhesion molecules, cytokine-cytokine receptor interaction, cholesterol metabolism, lipid, and atherosclerosis were common between the cord cervical and the spinal cord lumbar.

**Figure 8:**
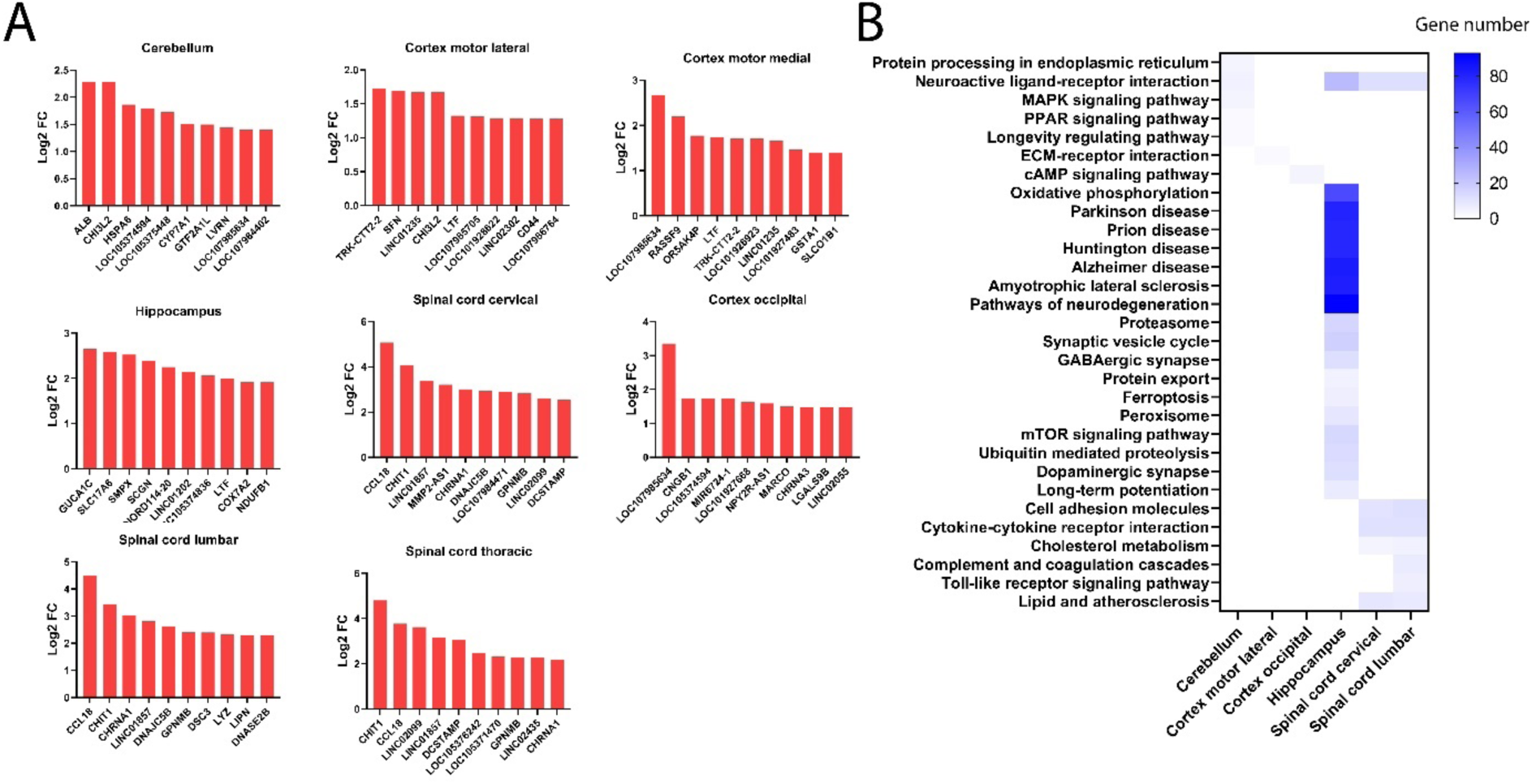
Upregulated and cellular pathways in ALS tissues. **A)** The bar plots show the top 10 upregulated genes in the cerebellum, lateral motor cortex, medial motor cortex, hippocampus, cervical spinal cord, and thoracic spinal cord of ALS patients. **B)** Pathways enrichment for upregulated genes (FC ≤ 2). In the hippocampus of ALS patients, pathways of neurodegeneration, ALS, Alzheimer’s, Parkinson’s, and Huntington’s diseases, were significant. Neuroactive ligand-receptor interaction was common among the hippocampus, cervical, and lumbar spinal cords.

#### 4.5.1. Gene expression of host genes and proteins that interact with ALS-related circRNAs

The expression of host genes of ALS-related circRNAs was verified in different tissues of ALS patients. According to the result, most of the host genes of these circRNAs are expressed in the tissues of ALS patients, especially phosphatidylinositol binding clathrin assembly protein (PICALM), chromosome segregation 1 like (CSE1L), transmembrane protein 38B (TMEM38B), exportin 1 (XPO1), and mitogen-activated protein kinase 1 (MAPK1) (Figure 9A). The expression of hub proteins was verified in the RNA-seq data from patients with ALS (Figure 9B). According to results, FMR1, ELAVL1, TIAL1, EIF4A3, and derine/arginine-rich splicing factor 1 (SRSF1) are significantly expressed in different tissues of ALS patients. Based on the DisGeNET database, only FUS is an ALS-associated protein (it is not shown in the figures).

**Figure 9.**
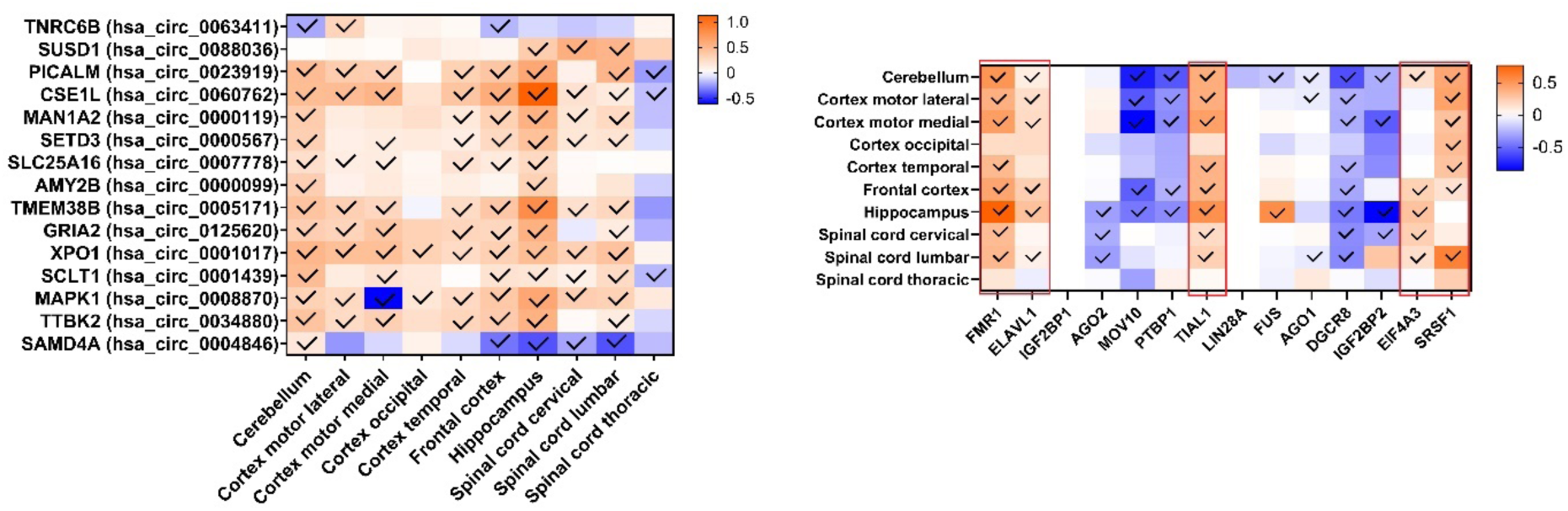
The expression of host genes of the ALS-related circRNAs and hub proteins that interact with ALS-related circRNAs. **A)** The expression of the host genes in different ALS patients’ tissues. Host genes, including PICALM, CSE1L, TMEM38B, XPO1, and MAPK1 upregulated in different tissues of ALS patients. **B)** Five genes of hub proteins FMR1, ELAVEL1, TIAL1, EIF4A3, and SRSF1 are significantly upregulated in tissues of ALS patients.

## Discussion

ALS is marked by progressive muscle weakness and paralysis, resulting from the deterioration of motor neurons. Our approach involved a comprehensive review of ALS-related circRNAs, followed by their analysis using bioinformatic tools. For this analysis, we exclusively focused on circRNAs with a circBase ID, as this ensured access to their essential sequence and fundamental data, which are critical for robust bioinformatic investigation, to elucidate the potential functions of ALS-related circRNAs.

According to the bioinformatic analysis, has_circ_0001017, has_circ_004846, has_circ_0034880, and especially has_circ_0000099, can rescue ALS-related genes more than the others. Our bioinformatic results showed that has_circ_0000099 can regulate different ALS-related cellular pathways, including complement and coagulation cascades, cytokine-cytokine receptor interaction, calcium signaling pathway, long-term potentiation, neuroactive ligand-receptor interaction, and PPAR signaling pathway through sponging miRNAs (Figure 3). Abnormal complement activity has been observed near degenerating motor neurons as ALS progresses. This, along with evidence of complement activation in the blood and muscle tissue of ALS patients, and an unexplained phenomenon of red blood cell toxicity in their plasma, strongly suggests that the complement system plays a significant role in how ALS develops and worsens (62). A distinguishing characteristic of MNs affected by ALS is their inherent vulnerability to calcium (Ca2+) overload. These neurons naturally express high levels of Ca2+-permeable AMPA receptors but have a low Ca2+ buffering capacity due to limited expression of intrinsic Ca2+ buffering proteins (CaBPs) like parvalbumin and calbindin. This combination of physiological features, while essential for normal function, likely predisposes MNs to the systemic intracellular Ca2+ overload observed in ALS. Interestingly, levels of calretinin and parvalbumin further decrease in the axons of MNs in ALS patients, exacerbating this buffering deficit under pathological conditions (63). In ALS, a neurodegenerative disease characterized by the progressive degeneration of motor neurons, the pathway of PPAR gamma is downregulated, while the canonical Wnt/beta-catenin pathway is upregulated. Despite this, studies on ALS transgenic mice have shown that treatment with PPAR gamma agonists can have beneficial effects, partly due to their anti-inflammatory properties. This suggests that modulating PPAR gamma could be a potential therapeutic strategy for ALS (64).

Moreover, pathway/biological process analysis for common gene targets of hsa_circ_0000099 showed that it can regulate some crucial cellular pathways in ALS, encompassing unfolded protein response, oxidative stress, and apoptosis (Figure 4). Dysfunction in protein homeostasis (proteostasis) is a key feature of ALS. The unfolded protein response and other proteostasis pathways are activated in both the motor cortex and spinal cord of sporadic ALS patients. In the motor cortex, specific unfolded protein response genes like PDIs are upregulated, which correlates with oligodendrocyte markers. In contrast, endoplasmic reticulum-associated degradation (ERAD) and heat shock response genes are predominantly activated in the spinal cord, correlating with neuronal markers. These findings indicate that while proteostasis disruption is a common mechanism in ALS, the specific cellular contributions differ between affected brain regions (65). In neurodegenerative diseases, such as ALS, oxidative stress and neuroinflammation are primary drivers of MN death. A central part of this process is mitochondrial damage, which can be detected early on. Therefore, effective treatment strategies should focus on targeting patients at or after symptom onset (66). The association of hsa_circ_0000099 in cell death and apoptosis has been studied in different experiments. Studies have shown that hsa_circ_0000099 regulates proliferation, invasion, and migration in SRA01/04 cells, which are treated with TGF-β2 through sponging miR-223-3p (67). However, this circRNA is downregulated in esophageal squamous cell carcinoma (68). hsa_circ_0000099 is also significantly downregulated in esophageal squamous cell carcinoma (69).

In the next step, the circRNA-protein interaction analysis was performed for the ALS-related circRNAs (Figure 6). Among the 20 obtained proteins, 7 of them interact with FUS. This interaction is significantly changed by FUS mutation hsa_circ_0000567 and hsa_circ_0060762 (Figure 7). The mutant FUS protein preferentially binds to specific introns that contain inverted Alu repeats, flanking the downregulated circRNAs. This suggests that the P525L FUS mutation disrupts the biogenesis and localization of circRNAs, which may contribute to ALS pathogenesis by acting as miRNA sponges (70).

we also employed the RNA-seq data to reach a gene expression profile of ALS (Figure 8). Most host genes of ALS-related circRNAs are expressed in different tissues of ALS patients. These changing expressions in host genes may correspond to changes in the expression of their circRNAs (Figure 9A). Therefore, they can be proper biomarkers for tissue impairment in ALS patients (71). Some host genes, such as AMY2B (host gene of hsa_circ_0000099) upregulated in just two tissues of ALS patients. This phenomenon might be a more specific sign of tissue impairment. Moreover, they are highly stable and abundant in the brain, which suggests they can pass into the CSF or blood if the blood-brain barrier is compromised. This makes them measurable in biofluids (72). Despite being similar in size to mRNAs, circRNAs are far more stable. CircRNAs can last for several days, whereas linear mRNAs typically degrade in less than 20 hours (73).

In addition, the gene expression of hub proteins that interact with ALS-related circRNAs was verified in the RNA-seq data. Results showed that FMR1, ELAVL1, TIAL1, EIF4A3, and SRSF1 were upregulated in tissues of ALS patients. The proteins UPF3B, regulator of nonsense mediated mRNA decay (UPF3B), and EIF4A3, which are components of the exon junction complex (EJC), were found to have increased levels in cell lines affected by ALS. Additionally, these proteins were observed to interact with FUS (74). SRSF1 is linked to ALS, playing a role in the nuclear export of pathological C9ORF72-ALS repeat transcripts. In this study, partial depletion of SRSF1 was investigated as a gene therapy approach. Results indicated that this depletion was sufficient to confer neuroprotection without globally disrupting gene expression, inducing specific beneficial RNA expression changes in ALS neurons and animal models (75).

## Conclusion

MNDs contain five main diseases: ALS, PLS, PBP, SMA, and PMA, which circRNAs dynamically active and associated with them. These circRNAs can regulate a wide range of miRNAs, genes, and cellular pathways. Regarding to bioinformatic analysis, among ALS-related circRNAs, hsa_circ_0000099, hsa_circ_0036411, hsa_circ_000088036, and hsa_circ_0060762 can regulate ALS-related cellular pathways including inflammation, cell cycle and apoptosis. Therefore, they can be potential targets for further studies and therapy of ALS.

